# Neuron-gated silicon nanowire field effect transistors to follow single spike propagation within neuronal network

**DOI:** 10.1101/2020.11.06.371369

**Authors:** C. Delacour, F. Veliev, T. Crozes, G. Bres, J. Minet, G. Becq, I. Ionica, T. Ernst, A. Briançon-Marjollet, M. Albrieux, C. Villard

## Abstract

Silicon nanowire field effect transistors SiNW-FETs provide a local probe for sensing neuronal activity at the subcellular scale, thanks to their nanometer size and ultrahigh sensitivity. The combination with micro-patterning or microfluidic techniques to build model neurons networks above SiNW arrays could allow monitoring spike propagation and tailor specific stimulations, being useful to investigate network communications at multiple scales, such as plasticity or computing processes. This versatile device could be useful in many research areas, including diagnosis, prosthesis, and health security. Using top-down silicon nanowires-based array, we show here the ability to record electrical signals from matured neurons with top-down silicon nanowires, such as local field potential and unitary spike within ex-vivo preparations and hippocampal neurons grown on chip respectively. Furthermore, we demonstrate the ability to guide neurites above the sensors array during 3 weeks of cultures and follow propagation of spikes along cells. Silicon nanowire field effect transistors are obtained by top-down approach with CMOS compatible technology, showing the possibility to implement them at manufacturing level. These results confirm further the potentiality of the approach to follow spike propagation over large distances and at precise location along neuronal cells, by providing a multiscale addressing at the nano and mesoscales.

## INTRODUCTION

During the last decades, the possibility to establish a bidirectional communication link with electrically excitable cells, such as neurons, becomes very attractive for both fundamental and medical purposes. Many conceptual and technological advances were achieved, including in the field of neuroelectronic, which remains a unique way to obtain quantitative measurements of neuronal electrical activity and tailor specific stimulations. Besides recording the activity of many neurons individually, electrical devices are indeed able to stimulate or inhibit neural spikes at single cell level within large networks, which makes them suitable tools to interrogate neural networks within the brain,^1^ slices or cell cultures.^2^

With nearly 40 years of development in that field, numerous materials and designs have been implemented on many substrates including transparent or flexible planar and 3D electrodes,^3–6^ but still the current conventional microelectrodes (MEs) face limitations to reach the ultimate sensitivity and to probe sub-cellular event such as ion channel, dendrite or synaptic activity that occurs at the nano and micro-scales.^7^ Their sensitivity and noise being proportional to the surface area, the MEs cannot be reduced below the cell size, being usually around 20 - 50 μm diameter wide for extracellular recording. Thus, MEs remain blind to the huge amount of information (weak signals) hidden within the dendrites, synapses, or ion channels. These key building blocks sustain all the brain processing and are involved in many neurodegenerative diseases, which motivates the development of nanoscale sensors, such as silicon nanowire field effect transistors (FETs). ^8–11^

Since the performance of FETs depends merely on their width-to-length ratio, the spatial resolution of the recording can be increased to the sub-micron range. In particular, the width of silicon nanowires could be reduced down to 6 to 10 nm, leading to an immense improvement of sensing properties for a variety of applications including real-time pH detection, chemical and bio-sensing.^12^ Additionally, the surface of SiNWs can be functionalized with catcher molecules such as antibodies,^13^ cancer markers,^14^ single-stranded DNA molecules^15,16^ and hybridization^17^ or virus^18^ to enable a specific detection of analytes for medical applications.^19,20^

Fromherz et al. was one of the first groups to record electrical signals from cells with silicon transistors, firstly on leach neurons in 1991,^21^ then on cultured cell lines such as cardiomyocyte^22^ or HL-1 cells^23,24^ and finally on isolated mammalian neurons.^25^ At the same time, Lieber’s group demonstrated the possibility to use ultrasensitive bottom-up silicon nanowire to record spikes from young cortical neurons (2-5 days old) deposited onto the nanosensors with PDMS stamp.^26^ These pioneer experiments have paved the road for further implementation of SiNW-FET at the manufacturing scale, to provide prototypes for various bio-applications, including cancer diagnostic, or detection of DNA, virus, bacteria and biotoxin, explosive gas, protein, nuclei acid for health and security.^27^

Here, we have fabricated defined arrays of (30-100 nm wide) silicon nanowire FET using top-down approach compatible with CMOS technology, and applied them to monitor electrical activity of several neuron assemblies: first within ex-vivo preparation (brain slice) deposited on the sample for the recording and secondly within hippocampal neurons cultured directly on the sample and maintained during several weeks. These devices enabled real-time monitoring of local field potential within the brain slices, as well as single spike from the matured neurons cultured on chip. We show the ability to guide the growth of neurites above the nanosensors, that allows a precise control of the neurons-nanowire positioning (even after 21 days in culture) and to follow the propagation of spike along single cell. Our results confirm the ability to detect locally electrical signals generated by the activity of neuron assemblies, as well as from individual neuron grown on chip, with top-down fabricated silicon nanowires. These results further demonstrate the potential of this scalable technology that could be implemented at manufacturing level (300mm Silicon-on-Insulator, SOI) and being useful to probe neuronal architectures at the meso and nanoscales.

## MATERIALS AND METHODS

### Fabrication of silicon nanowire FET arrays

Starting from high quality SOI substrates (SOITEC, France), single crystalline top Si-layer were selectively doped with boron atoms to form low doped (p-type, or pristine) nanowires with highly conductive (p++) drain-source contact areas. First, the undoped SOI substrate is oxidized to thin down the Si top layer to the desired thickness (30 - 50 nm), which is then doped by plasma assisted (Boron) ion implantation technique. Next the ohmic contacts are defined by optical lithography using positive resist, followed by evaporation of metals (Ti/Pt/Au) and the resist lift-off. Using e-beam lithography, nanowires with different dimension and geometry can be realized on the same chip (figure 1a, S1). The width of the NWs hereby varies from 30 nm to 1 μm and the length from 6 to 10 μm. Typically 90-80 NWs per chip are exposed and etched using reactive ion plasma etching (figure 1b-c). The contact resistance is reduced by a factor 100, after a rapid thermal annealing at (1h, 500°C) within inert (Ar/H_2_) atmosphere. The annealing forms a conductive titanium silicide alloy, which lowers the Schottky barrier height between Ti and Si.^28^ Finally, a thin gate high-k dielectric material is deposited using atomic layer deposition (ALD) above the nanowires (5-10 nm HfO_2_), and the metallic leads are passivated with thick isolating layer (parylene-C or silica) to avoid any leakage current in liquid gate operation. The thick oxide is removed above the nanowires to expose them to the cells with photolithography and reactive ion plasma etching (figure 1d-e).

**FIGURE 1.**
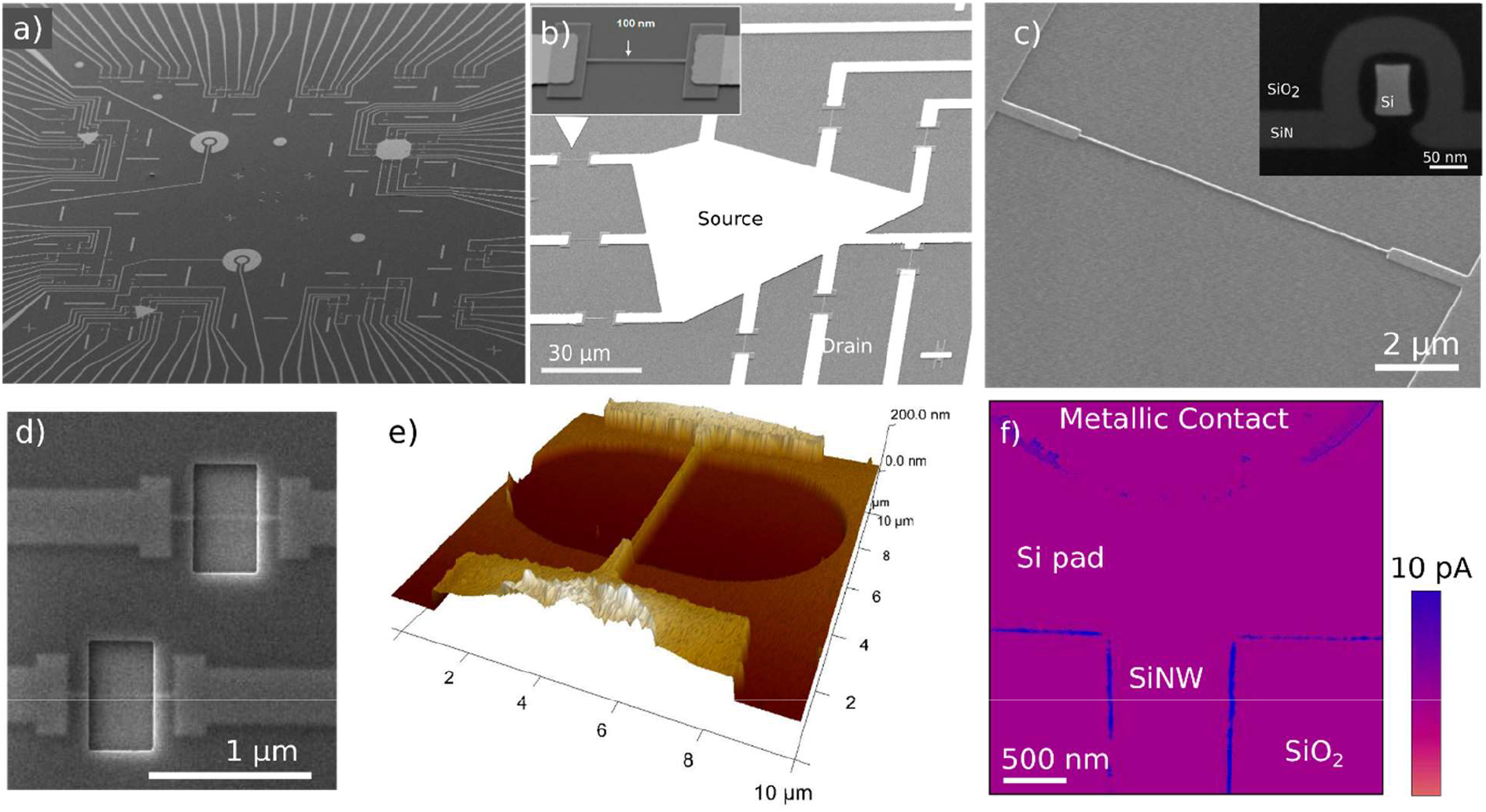
Silicon nanowire FETs fabrication. a-d) Representative scanning electron micrographs of the sample and device: a) several arrays of silicon nanowire transistors. The sensing area is about 10×10 mm^2^ and contained around 100 nanowires. b-c) Silicon nanowire (100 nm and 30 nm wide respectively) before the passivation of the metal contacts and cross-section of a nanowire embedded in a thick silicon oxide covering the metallic leads for operation in liquids (inset 1c). d-e) The SEM and atomic force micrographs show the final devices, after removing the thick oxide layer on top of the nanowire. f) TUNA Tunneling atomic force micrograph of a silicon nanowire covered by the thin top oxide (5 nm, HfO_2_) showing weak leakage current value with the front gate (few pA, V_bias_ = −2V). Nanowires reported figures 1c-d have been realized with CMOS clean standard process.

Silicon nanowire arrays have also been fabricated in CMOS technology platform on 300 mm SOI wafers. Similar process flow was used, except that the thin gate oxide (SiO_2_) is thermally grown above the nanowire. This few-nanometers-thick passivation layer is protected by a SiN coating when removing the thicker silica oxide (300 nm) above the nanowire that isolates the metallic pads (figure 1c inset). The SiN layer is finally removed for sensing operation (figure 1d).

**Electrical characterizations** of the nanowires are characterized under ambient conditions, using a shield multi-probe station (figures 2a, 3a) connected either to a Keithley source-meter or a FPGA (fast programmable gate array) card interfaced with LabView for polarization and acquisition. Te nanowires are voltage biased while a low noise femto-current amplifier monitors the drain source current. Input and output cables are filtered to remove all the high frequency (>2kHz) parasitic signals. Electrical setup and schematics are shown with the electrical characterizations (figures 2 and 3).

**FIGURE 2.**
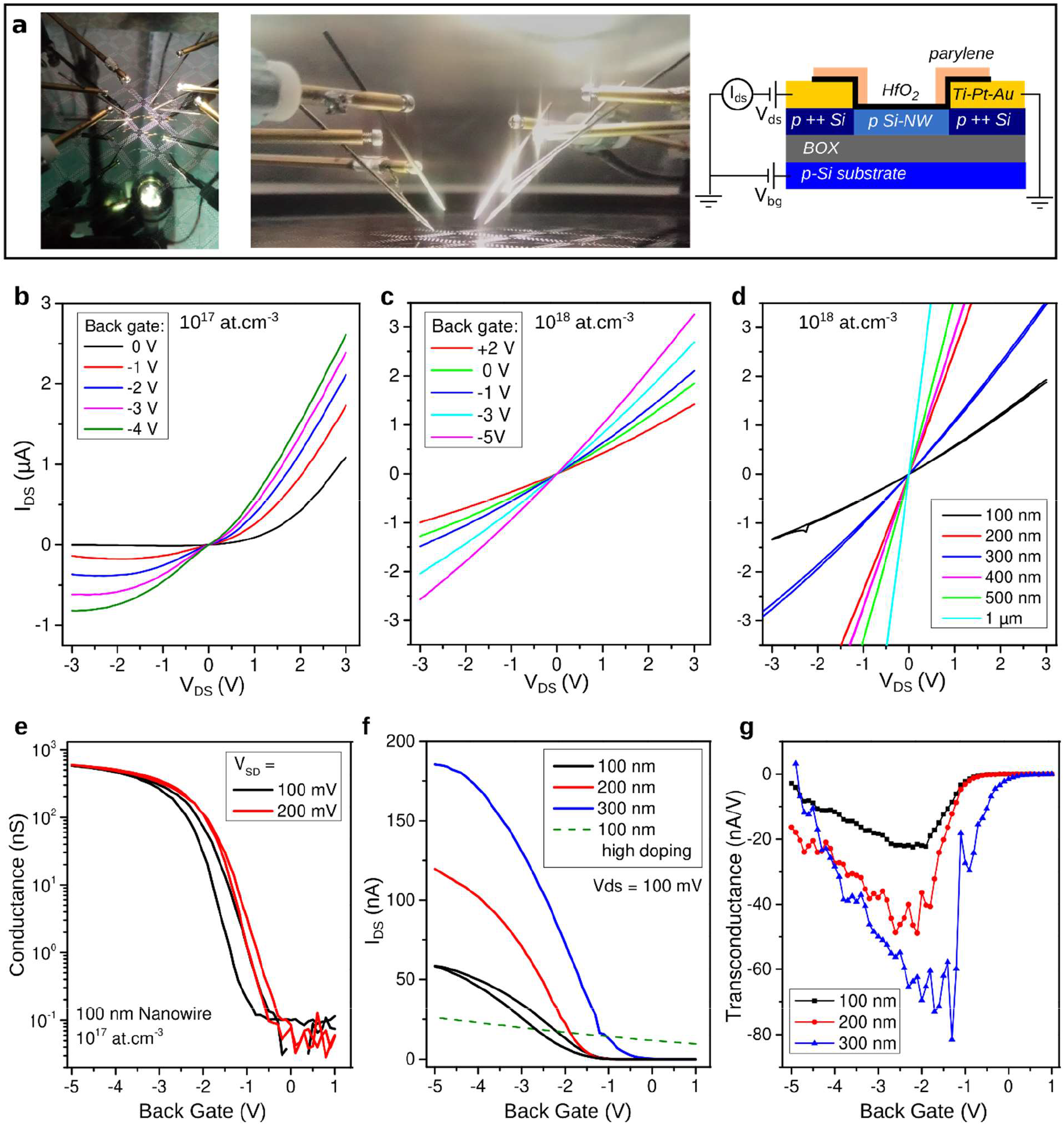
Back-gated silicon nanowire FETs. a) Pictures of the probe station and chip array integrated on 300mm wafer, and schematic of the measurement showing a cross section (out-plan) of the FETs. The conduction channel and leads are p-doped (10^17^ - 10^18^ at/cm^3^ and 10^21^ at/cm^3^ respectively). (b-c) I_DS_(V_DS_) curves of a 10^17^ at/cm^3^ and 10^18^ at/cm^3^ doped SiNW-FET for several back-gate voltage, and I_DS_(V_DS_) curves of SiNW-FETs with varying channel width. (e-g) Modulation of the silicon nanowire conductance when applying a back-gate voltage on the low p-doped Si substrate. The I_DS_(V_DS_) and I_DS_(V_BG_) curves are measured for several bias voltage (e) and SiNW-width respectively (f). g) Transconductance versus back gate voltage for nanowires with different widths at V_Ds_ = 100 mV.

**FIGURE 3:**
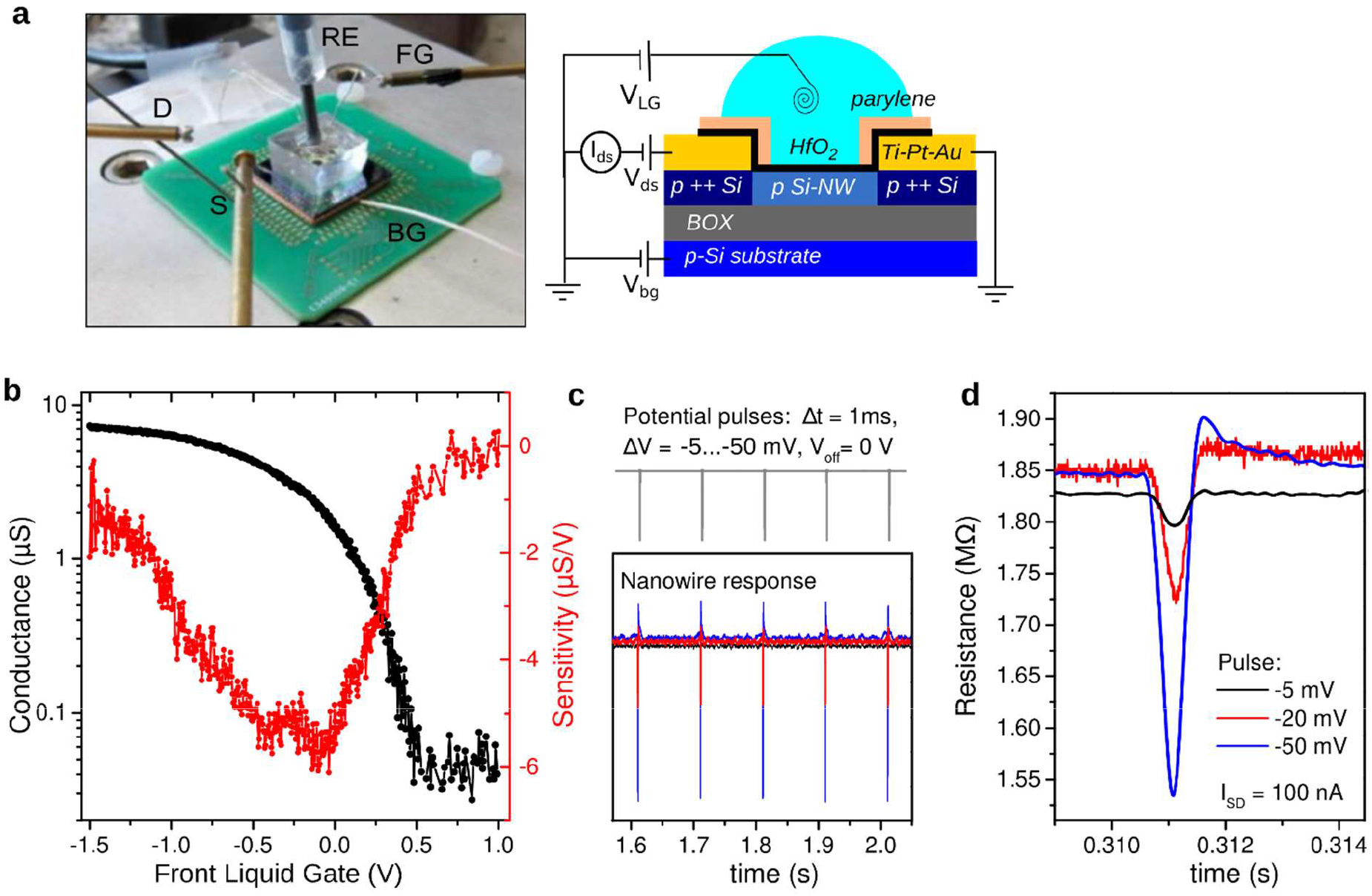
Liquid gated silicon nanowire FETs. a) Picture and schematic of the liquid gated operation. The drain-source current I_DS_ is monitored as function of the front (FG) and back (BG) gate voltages. The Ag/AgCl reference electrode (RE) is used in addition with a Pt wire (FG) immersed in the solution. b) Modulation of the conductance of a 100 nm wide NW-FET by varying the front liquid gate voltage through the Pt-electrode. c) Detection of short (Δτ=1 ms) potential pulses applied to the front liquid gate using SiNW-FET. d) Zoom of the potential pulses with different amplitudes detected by the SiNW-FET at bias current I_DS_ = 100 nA. The potential pulse induces a resistance/conductance change of the nanowire channel.

### Neuronal signal monitoring

A complete, portable and reliable system has been developed to monitor the channel resistance of 12 transistors simultaneously at 10kHz sampling frequency each and fixed current I_DS_ (figure 5a and 5b). A bipolar modulation of the bias current is used to cancel all parasitic electromagnetic (e.m.f.) signals. The current source ranges from 200nA to 700nA, that could be changed by a polarization resistor switch. All acquisition parameters (sample frequency, current setpoint, biased transistors) can be changed in real-time, with an intuitive software coded in Delphi. Analog to digital converter ADC readings are synchronized by the hardware. The system, composed by the printed circuit board PCB and the software is able to display simultaneously 12 transistors impedances variation.

### Acute brain slices

Sagittal slices containing the rostral midbrain (300 μm thick) were prepared from young male Sprague-Dawley rats (postnatal days 17-21; Janvier, Le Genest St Isle, France). All experiments were performed according to animal care guidelines in agreement with the European Community Council Directive of November 24, 1986 (86/609/EEC), in accordance with the French guidelines on the use of living animals in scientific investigations and with the approval of the “Grenoble Institute of Neurosciences ethical committee”. Rats were killed by decapitation and their brains were removed rapidly and cut into slices with a vibratome VT1000S (Leica, Wetzlar, Germany) in cold low Ca^2+^-high Mg^2+^ artificial cerebral spinal fluid (ACSF) containing: 126 mM NaCl, 2.5 mM KCl, 7 mM MgCl_2_, 0.5 mM CaCl_2_, 1.2 mM NaH_2_PO_4_, 25 mM NaHCO_3_, 11 mM D-glucose, bubbled with 95% O_2_/5% CO_2_. Sagittal slices containing the Substantia Nigra pars reticulata (SNr) were placed in ACSF (126 mM NaCl, 2.5 mM KCl, 1.2 mM MgSO_4_, 2.5 mM CaCl_2_, 1.2 mM NaH_2_PO_4_, 25 mM NaHCO_3_, 11 mM D-glucose) saturated with 95% O_2_/5% CO_2_ and supplemented with 1 mM sodium pyruvate at room temperature, for a recovery period.

### Primary hippocampal neurons culture

E16.5 pregnant mice were killed by cervical elongation. Hippocampi were dissected from the mouse embryos in HBSS-HEPES (Gibco), incubated 15mins in Trypsin 2,5% (Gibco), rinsed in HBSS-HEPES twice, and mechanically dissociated by repeatedly pipetting in 1ml attachment medium (MEM supplemented with 10% foetal bovine serum, 1% glutamine and 0.05% peniciline/streptomycine, Gibco). Cells were seeded with a density of 12 000 cells/cm^2^ on the chip previously sterilized and coated with poly-L-lysine (PLL, 1mg/ml) to ensure neurons adhesion. PLL micropatterns are performed by photolithography before the culture, to guide the neurites outgrowth along defined pathway (figure S1). The neurochips were then incubated at 37°C and 5% CO_2_ in the attachment medium, and replaced a few hours later by glial conditioned medium supplemented with 1 mM AraC. To obtain glial conditioned medium, glial cells from E16.5 mouse embryo cortex, were cultured in MEMc +10% FBS until confluency. The cells were then cultivated with serum free Neurobasal medium (Invitrogen) and the medium was collected after 48 h for subsequent use on neuron cultures. The culture medium was changed every 7 days, by replacing one-third of the volume with fresh astrocyte conditioned medium. The growth on silicon chip was observed using upright optical microscope (Olympus X51) and with conventional transmission microscope on the control glass coverslips.

### Immunostaining

After the recording neurons are fixed in 3.9 % paraformaldehyde (10 min), permeabilized in PBS-0.25% Triton-X100, and blocked in PBS-2% bovine serum albumin. Staining is then performed using the anti-Synapsin (1:500, Millipore), anti-YL1/2 (1:1000, BioRad) primary antibodies, and DAPI (1:1000, Sigma-Aldrich) for labeling the synaptic vesicles, the cytoskeleton and the nucleus respectively. Cells were washed in PBS and incubated with FITC and TRITC-conjugated secondary antibodies (Alexa Fluor, dilution 1/500). Cells were washed again in PBS and mounted in Mowiol (Sigma-Aldrich) mounting medium. Immunofluorescence images were collected using an Olympus BX51 and immersive objectives x20 or x60.

## RESULTS AND DISCUSSION

### Array of silicon nanowire FETs

There are two main approaches for the fabrication of silicon nanowire, the bottom-up synthesis (typically by chemical vapor deposition CVD or molecular beam epitaxy) or the top-down nano-structuration (silicon etching or oxidation) of single crystalline silicon-on-insulator SOI wafer. While bottom-up fabrication leads to narrowest nanowires and highest mobility values,^29,30^ the main drawback is to organize the nanowires at the mesoscale to obtain FET arrays with defined geometry and reliable electrical properties. Here, we have rather used the counterpart top-down approach, which is compatible with CMOS technology enabling large scale implementation of the nanowires with excellent and reliable electronic properties, and which provides an accurate positioning of the nanowires (fabrication process is detailed in methods).

Examples of the sample and nanodevice array are shown within figure 1. The arrays of nanowires can be designed to interface simple models of neurons networks, such as three-nodes closed loop network (figure 1a-b and S1).^31^ The SEM micrographs demonstrated the overall homogeneity and high quality of the silicon nanowires obtained with e-beam lithography and silicon etching. Figures 1c and 1d are representatives of silicon nanowires fabricated within CMOS technology platform on 300 mm SOI wafers, that enable to further reduce their size down to 30 nm wide in a reliable manner even for long nanowires (typ. 6-10μm).

The insulating properties of the thin high-k dielectric material deposited above the nanowire (typ. 5 nm thick HfO_2_) have been further investigated using tunneling atomic force microscopy (figure 1f). The leakage current from the device to the metallic AFM tip is well homogeneous over the sample surface, being around few pA (at 1V bias voltage) in agreement with gate source current *I*_*GS*_ measured with the probe station. Higher values appear on sidewalls of the nanowire and metallic pads that certainly stem from a larger contact area with the AFM tip. This result demonstrates the homogeneous coating and efficiency of the top oxide above the nanowires.

### Electronic properties

As described in figure 2, the drain-source current through the transistor channel I_DS_ is measured at varying polarization V_DS_ and back-gate voltage V_G_. As expected for the low doped p-type nanowires 10^17^ at/cm^3^, I_DS_ saturates with increasing drain source voltage (negative V_DS_) due to narrowing the FET conduction channel at the drain contact (figure 2b). We also fabricated highly doped nanowires (10^18^-10^19^ at/cm^3^), but the devices exhibit a rather ohmic behavior at low drain source voltage (figure 2c). As expected, the net current through the nanowire increases almost linearly hereby with its width (figure 2d).

When applying negative back-gate voltages, the majority carriers (holes) are accumulated, increasing the conductivity of the nanowire channel until the free carriers are exploited entirely and the conductance C_DS_ saturates (figure 2e). For more positive V_G_ values the nanowire is depleted, and no current is flowing through the channel due to the forming of p-n junctions between the depleted nanowire and heavily p-doped contact regions on source and drain sides. The back-gate induced conductance modulation is stronger for low doped SiNWs, and the current through the channel scales with the width of the nanowire (figure 2f). As expected the sensitivity, proportional to the normalized transconductance 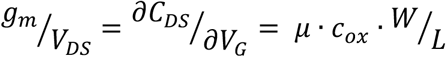 scales with the width-to-length ratio W/L (figure 2g).

The effective carrier mobility can be extracted from the transfer curve in the linear regime according to the previous expression. To estimate the interfacial capacitance, we consider the nanowire as a wire of radius R embedded in a dielectric *ε*_r_ on an infinite conductive plate in distance d (200nm of SOI-BOX oxide). The analytic expression of the gate capacitance is then given by^32^:

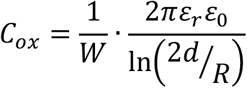

with less than 1% error for d/R > 6. In our case, however the nanowire is lying on the silicon dioxide (back gate), which leads to a smaller effective dielectric constant: 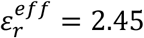. The effective radius R of the nanowire can be defined as the radius of a circle with the same area as the nanowire cross section. For the trapezoidal nanowires with widths of 100 nm and 50 nm high, the effective radius is 37.8 nm and the gate capacitance is c_ox_ = 5.6.10^−4^ F/m^2^. With a transconductance of g_m_=20 nA.V^−1^ at V_D_ = 100 mV (figure 2g), this leads to an effective hole mobility μ_h_ = 250 cm^2^.V^− 1^.s^− 1^, which is in the range of the bulk values with similar doping concentration, and slightly higher than previously reported value for top-down fabricated nanowires.^30,33,34^

The liquid gating properties of the SINWs are tested in water, saline solution and cell culture medium (Neurobasal, Invitrogen), by monitoring the nanowires conductance while sweeping the front gate potential with a quasi-reference Pt electrode immersed in the electrolyte above the nanowires. At the same time, a silver chloride Ag/AgCl reference electrode is used to measure the liquid potential as shown figure 3a. In analogy to the back-gate, the conductance increases due to the charge carrier accumulation by applying more negative voltages to the liquid gate. The maximum sensitivity is around 6 μS/V, considerably higher than with a back-gate 0.2 μS/V due to the thinner top gate oxide (few nms) and thus higher gate capacitance (figure 3b).

We have assessed the ability of silicon nanowire to detect short pulses similar to action potentials in neurons. Gaussian potential pulses (1 ms long) with varying amplitude *V*_*p*_ were applied to the cell culture medium through the Pt-gate electrode, while the voltage drop at the nanowire was measured at fixed polarization current I_DS_ = 100 nA. The DC-offset of the gate electrode was kept at 0 V to ensure the highest sensitivity (figure 3b). The potential pulses detected by nanowires can be seen in figure 3c-d. The detected signal exhibits slightly biphasic shape, indicating that the field effect induced conductance modulation (monophasic negative pulse: *I*_*DS*_ = *g*_*m*_ · *V*_*p*_) is superimposed with a slight capacitive current due to the rapid potential change (*I*_*c*_ = *C* · *∂V*_*p*_/*∂t*). The signal to noise ratio remains high (>3) even for weaker pulses (−5mV) enabling an efficient spike detection.

### Ions sensitive field effect transistors

We have assessed the pH sensitivity of the nanowires. For that, the fluidic chamber above the nanowires is subsequently filled with a solution of different hydrogen ion concentration, while the FET conductance is monitored. With increasing pH, we observe an increase of the transistor channel current I_DS_. The depletion of positively charged hydrogen ions leads to more negatively charge surface at the oxide/solution interface. As a result, the concentration of majority carrier increases within the nanowire conduction channel and so does the drain-source current. This pH-induced current modulation is fully reversible and relatively stable in time (figure 4b).

**FIGURE 4.**
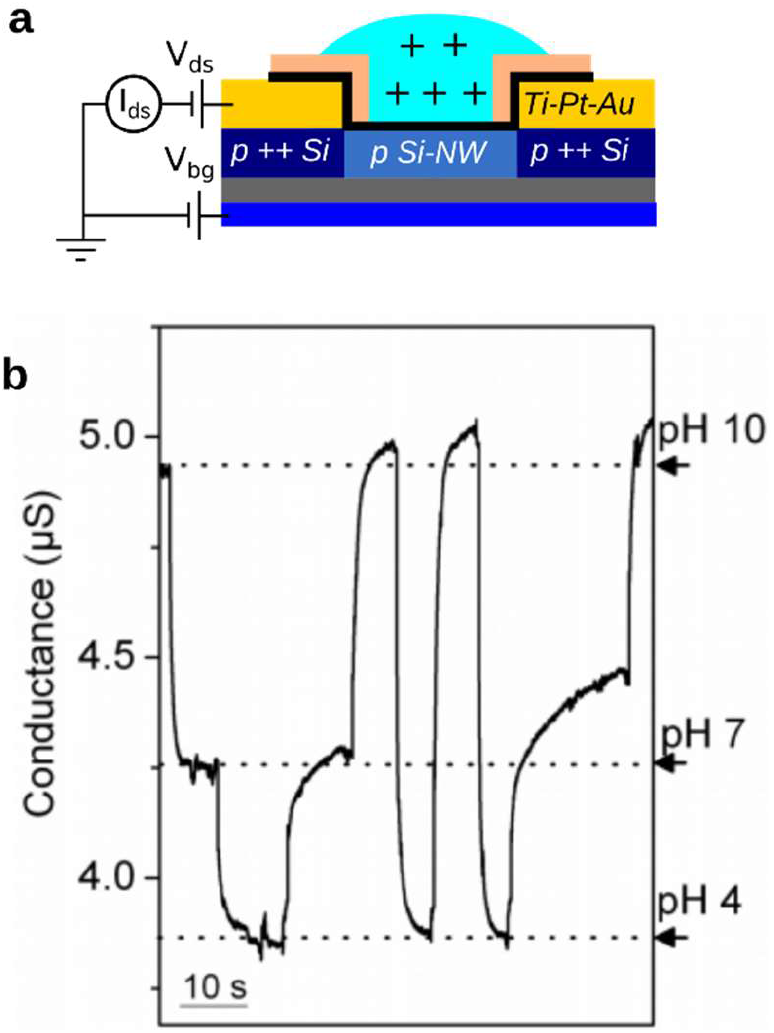
pH detection with silicon nanowire FETs. a) Schematic of the pH detection. The drain-source current is recorded while changing the electrolyte with solutions of several pH values. The concentration of positive charges at the oxide/solution interface varies and modulates the concentration of majority carrier within the transistor channel. b) Detection of pH changes using a SiNW-FET (100 nm wide, 10^17^ at.cm^−3^) at the reference liquid gate potential set to 0 V.

The pH-sensitivity of the nanowires can be estimated according to the following expression:^35^

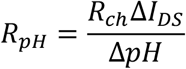

with R_ch_ the transistor channel resistance and Δ*I*_*DS*_ the modulation of current induced by a change of ΔpH. The sensitivity of the nanowire is 46.4 mV/pH (at zero back gate voltage and V_DS_=1V) in agreement with the Nernst limit 59.5mV/pH for an ideal surface.^36^

The devices sensitivity are almost one order of magnitude lower than bottom-up SiNW-FETs in electron regime, that reach mobility value of about 1350 cm^2^.V^− 1^.s^− 1^.^29^ However, these nanowires exhibit similar performance compared to the top-down SiNW-FETs which have been used for biosensing and for interfacing electrogenic cells,^23,37^ ie 2-6 μS/V and 41mV/pH. Also, The application of a negative back gate voltage could be used to further improve the pH sensitivity of the nanowires.^35^

### Brain slices recording

The silicon nanowire arrays were interfaced with acute rat brain slices to assess their ability to record local field potential generated by the collective activity of a neuron assembly. Specially, we targeted the substantia nigra (pars reticulata, SNr) which is a major output nucleus of the basal ganglia circuitry located in the midbrain, mostly studied for motor disorders characteristic of Parkinson’s disease. Interestingly, these networks exhibit a periodic activity with a frequency of about 10 Hz, ^38,39^ which could facilitate the signal sorting analysis.

Isolated slices were placed onto the SiNW-chip in the tissue medium solution regulated at room temperature and bubbled with 95% O_2_, 5% CO_2_ (figure 5c) The slice position over the chip is then adjusted to align the SNr area with the SiNW arrays, and the conductance of the nanowires are monitored along time, at constant drain-source voltages using our home-made electronics (see materials and methods, figure 5a-b).

**FIGURE 5.**
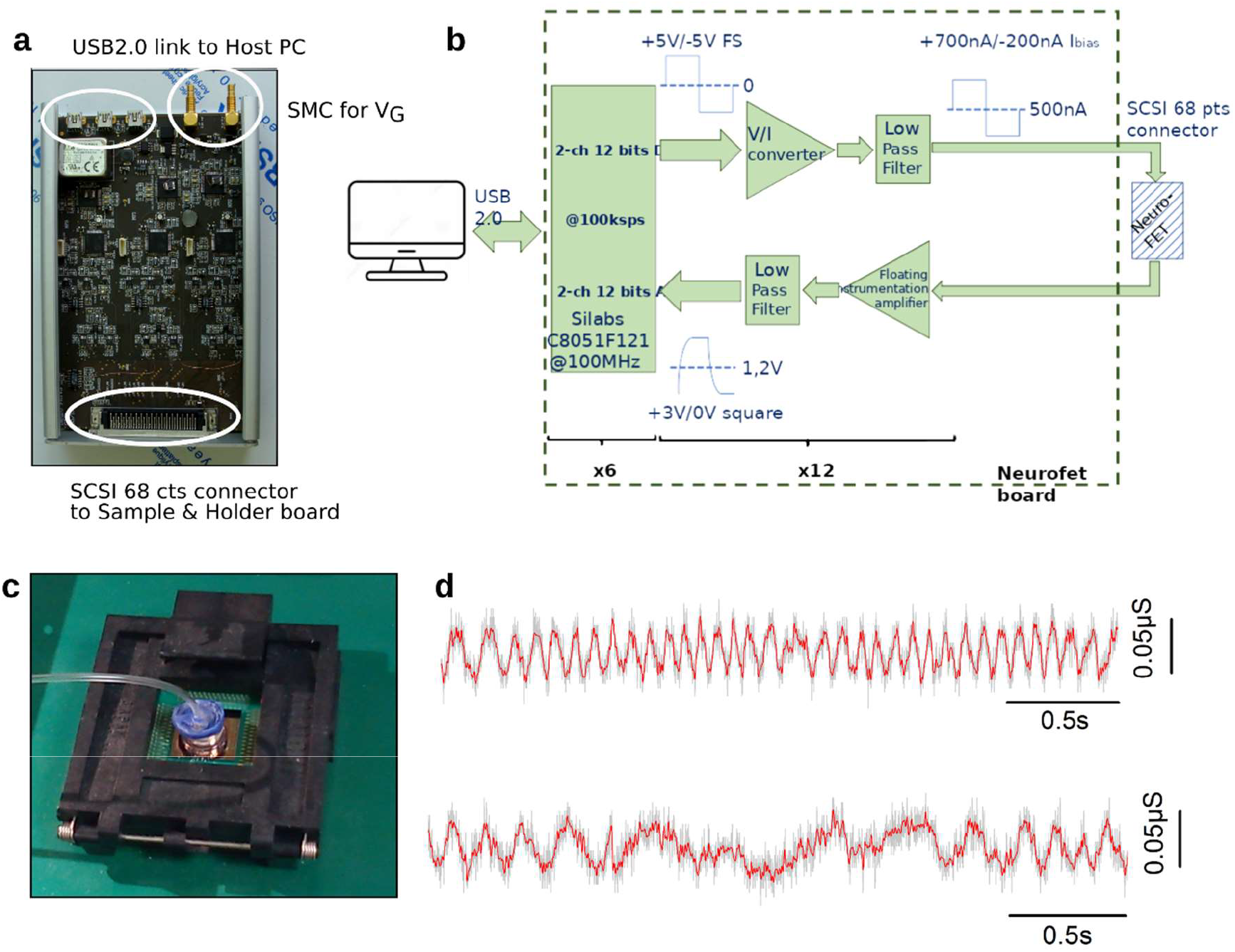
LFP recording with silicon nanowire FETs. a) Electronic card dedicated for electrophysiological recordings with the SiNW arrays. The card is connected to the PC via USB port and piloted with Delphi coded software. b) Principle of the electrical setup. NWs are biased with a low noise current source, while the drain-source voltage is amplified and low-pass filtered (12 nanowires simultaneously). c) Picture of the PCB-connector and silicon chip. A 12mm wide glass chamber (regulated in CO_2_) contains the recording solution and the brain slice from the Substantia Nigra pars reticula (SNr) of rat (detailed in methods). d) Typical time traces of the nanowire conductance at several time windows, from the beginning (upper) to the end (lower) of the recording. Local field potential (LFP) generated by the assembly of the SNr neurons induces slow oscillations (8-12 Hz, about 50 nS) of the transistor channel conductance.

Figure 5d shows a representative conductance time-trace recorded from the nanowires, and the expected low frequency oscillations characteristic of the SNr electrical activity. As the cells are still embedded in their highly connected network and matrix and relatively far from the sensor, the collected extracellular voltage rather results from the activity of many neurons than from a single spike/neuron. The frequency of the oscillation varies from 14 Hz to 7 Hz during the time of the recordings which is in good agreement with the frequency measured in living rats.^39^ The estimated extracellular potential is about 5 mV, using sensitivity value (6 μS/V) estimated previously (figure 3), which is slightly higher than the values reported *in-vivo* (around 1mV) measured with nickel-chrome microwires.

### Single spike detection within cultured neurons

Primary cultures of hippocampal neurons are performed in-situ on the silicon nanowire FET arrays until maturation (detailed in material and method). After 19-21 days in culture, the neurons and the network are mature in terms of electrical activity and connectivity. The synapses distribution is dense along the neurites as observed with immunofluorescent staining (figure 6a) as well as the network formed by the neurites that are both as expected for matured neurons. The overall activity of the network is confirmed on control sample by patch clamp measurements and calcium imaging (data not shown).

**FIGURE 6.**
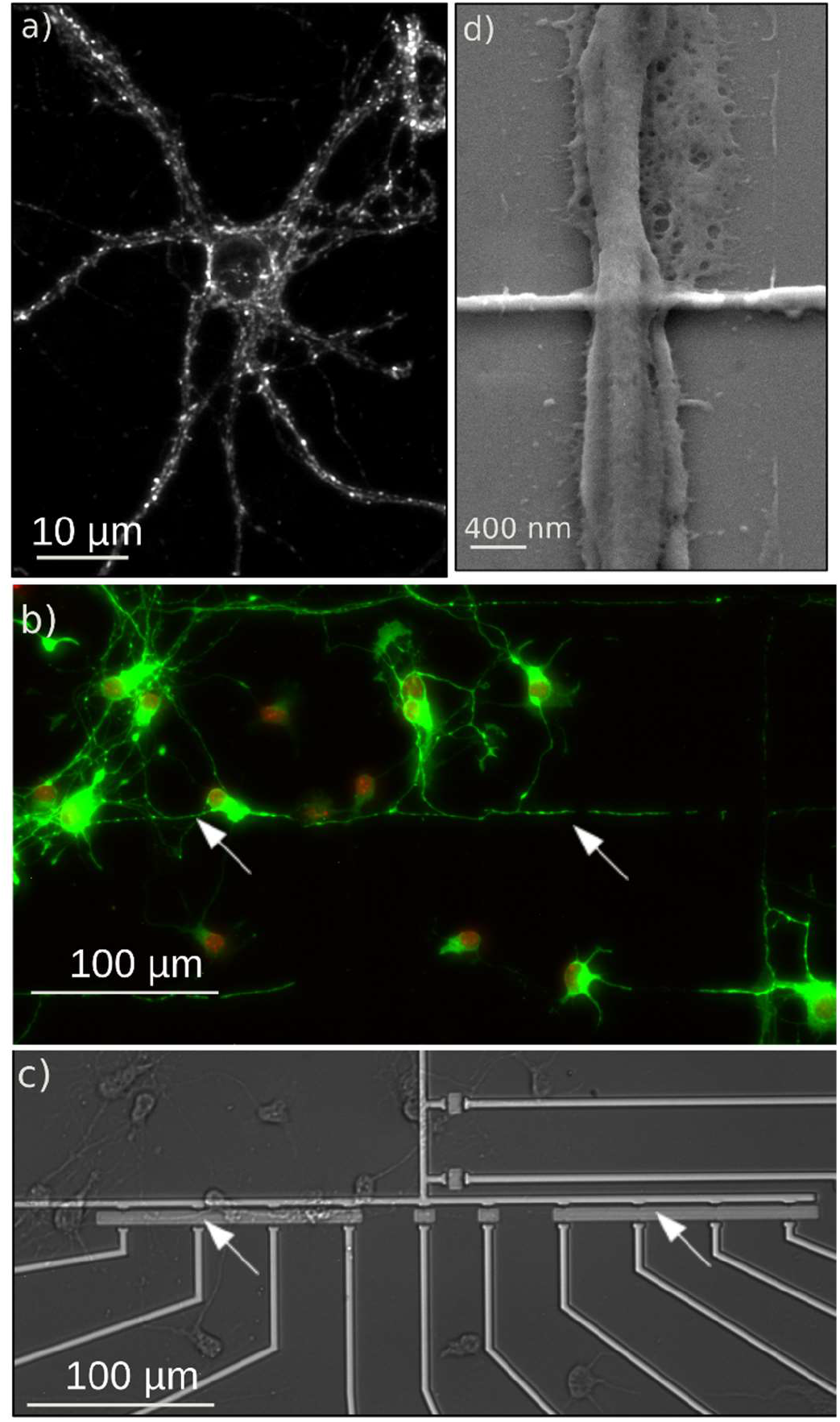
Neurons cultured and guided above silicon nanowires FETs. (a) Immuno-fluorescent (IF) micrograph (60X) of hippocampal neurons (21 DIV) labeled with anti-synapsin to show the distribution of synapses along the cell membrane (white dots). Fluorescent (b) and (c) optical micrographs of mature hippocampal neurons guided above the nanowire array with poly-L-lysin adhesive micropatterns (21 days of culture). The immunostaining (b) with anti-tubulin (green) and DAPI (red) show cell arborization and nucleus respectively. The arrows underline aligned neurites crossing several SiNWs. d) SEM micrograph providing a closer view of a neurite crossing with a nanowire. The nanowire is embedded by the plasma membrane.

Neurons are guided over the array of nanowires with patterns of poly-L-lysin molecules.^31^ An example is shown figure S1. The efficiency of the adhesive patterns is enough to keep several neurites above the sensors after 3 weeks in culture, even if neurons start to spread outside the patterns as shown within figure 6b-c. The cultured neurons strongly adhere on the substrate and its topography. As observed within figure 6d, the nanowires are well embedded by neurons which wrap the three faces of the nanowires. Such high covering of the nanowire by the neurons should also increase the sealing resistance and thus the device response as observed for mushroom-like structure,^40^ in comparison with planar microelectrode.

After 3 weeks of neurons culture, the conductance of the nanowires is monitored at fixed bias current, as described previously within materials and methods section. The addition of bicuculline (BIC, 20 μM, 15 min 37 °C) - a GABA_A_ receptor antagonist which activates spiking activity - in the culture medium allows to increase the probability to detect single spike as there are few neurons above the sensor array. The time traces of the conductance of two neighboring SiNW-FETs (40 μm apart) exhibit high spiking activity (figure 7a), with several short pulses (1-2ms) and longer bursts, that are characteristic of neuronal activity. The response of the nanowires is higher than for extracellular local field potential recorded from ex-vivo preparation (figure 5d). The polarity is also opposite. The conductance changes ΔC range from minus 100-200 nS, which correspond to a positive potential change of 16-33 mV with a typical nanowire sensitivity of −6 μS/V (figure 7b). The responses of the nanowires indicate that majority carriers are repulsed from the transistor channel, in result of an increase of positive charges at the top gate interface that could be caused by inward current of Na^+^ sodium or Ca^2+^ calcium ions during either action potential spike or post-synaptic currents propagation. These values suggest that the nanowires record intracellular potential changes, which can occur in case of a strong ohmic coupling to a leaky neuron membrane and when the nanostructure are engulfed by the plasma membrane. ^9,40–44^

**FIGURE 7.**
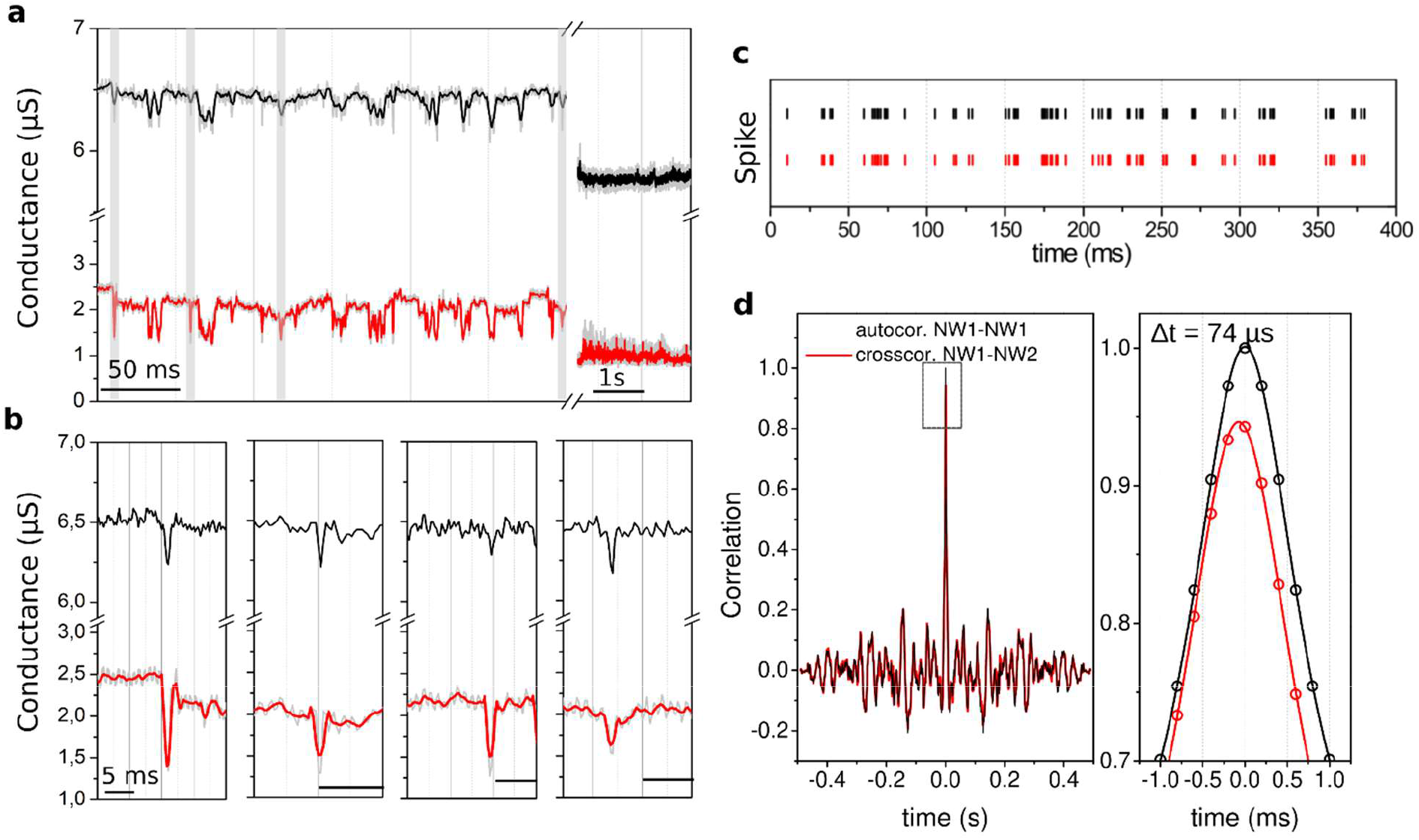
Neuron-gated silicon nanowire FETs. a) Conductance time-trace for two neighboring SiNW-FETs (39 μm apart) on which neurons have been cultured 21 days. Recording is performed in the culture medium with addition of bicuculine to increase the spiking activity. Without temperature and CO_2_ regulation, neurons died after about 30 mins of recording. At this stage (right time-windows, t >10000ms), spikes are suppressed and the nanowire conductance go back to its initial value (without neuron). (b) Zoomed view of representative spikes detected with the NWs, highlighted in (a). Neuronal spike induces a negative change of NW conductance of about minus 100-200 nS and duration of 1-2 ms. (c-d) Raster plots and cross-correlation of the nanowires conductance time traces shown in (a). The zoomed view (right panel) reveals a delay of about 74 μs between the two plots.

The traces detected on two adjacent nanowires are well correlated as shown in the raster plots (figure 7c). The cross-correlation between the two recording traces (for NW1 and NW2) shows a clearly identifiable shift, which corresponds to a time delay of Δτ = 74 μs (figure 7d). Same time delay was also found by comparing individually the shift between the spikes detected at each nanowire. For the conductance modulation due to the potential fluctuations in the cell culture medium or due to the electrical noise of the setup, the cross-correlation function would be centered at zero. The observed shift however suggests a propagating signal. The propagation velocity can be calculated to v = d/Δτ = 0.52 m/s, which is in good agreement with the values reported for propagation of neural spikes in cultured hippocampal neurons. ^26^

The conductance time traces of the nanowire were recorded after the experiment (right time windows in figure 7a). When neurons die, the short spikes are suppressed and the conductance of the nanowires decrease down to its initial value (without neurons), which is consistent with a disruption of the negatively charged plasma membrane.

### Discussion

Detection of neuronal activity on patterned neuronal networks is extremely challenging due to several reasons. Firstly, the neural maturation, i.e. the establishment of spontaneous electrical activity, is delayed on isolated networks containing only few neurons,^45^ while extended culturing periods usually result in soma displacement and clustering. Secondly, the contact area of the neurite with the FET device is very small, around 1/5 of the FET channel, which decreases the effective sensitivity. The potentially detected signal can be buried into the recording noise. Decreasing the NW length would improve the coupling, but also complicate the neural patterning.

The fabricated SiNW-FETs were able to detect local field potentials generated in acute brain slices, follow signals propagation within hippocampal neuron cultures and defined neuronal networks with an exact positioning of neurites above the transistors. The ability to keep neurons above the devices during 3 weeks seems favor an optimized coupling with the cell. The neurons may engulf the nanowire, providing high conductance modulation in comparison with previous planar Si- and SiNW-FETs,^22,23,25,46^ with recorded potential ranging from 100-200 μV up to 1 mV.^25^ This kind of recordings strongly depends on the transistor dimensions (which affect the sensitivity) and cell type interfaced to the FET. As reported in several studies, the coupling could change by several magnitude the response of neuron gated FET.^44^ Our results suggest that our system enables an optimized coupling, both on morphological and electrical point of views, between the FETs and the neurites.

## CONCLUSION

We have implemented arrays of three-gated (30nm thick) silicon nanowires with top down approach for biosensing applications. The fabrication of the nanosensors is compatible with CMOS technology for large scale implementation on 300mm substrate. The devices exhibit performance comparable with the state of the art and enable to detect a wide range of neuronal signal from low frequency LFP oscillatory waves to single spike within *in-vitro* mature networks. We have demonstrated the ability to guide neurite along the nanosensors arrays, opening the way to follow single spike propagation at the single cell level and along mesoscale network. Our measurements suggest high values of the recorded voltage at the cell-sensor junction, that reveal a strong coupling with the cell. This feature would be an advantage for improving detection efficiency, especially for weak neuronal signals along neuritis. Our work also combines a nanoscale recording localized at specific location along cell by using adhesive patterns to guide neurites above the nanosensors. Further investigations combining patch clamp with SiNW-FETs for instance would be useful to investigate further that interface and to decipher the cell-device coupling features in future works.

## Supplementary Information

### Supplementary figure

**Figure S1.**
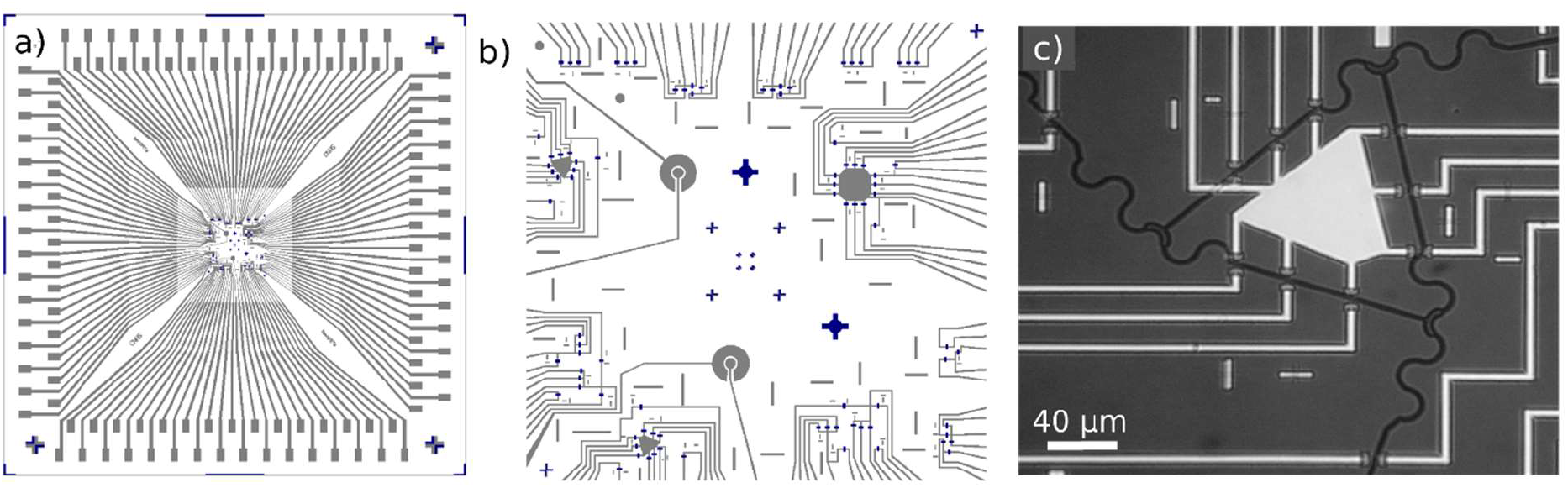
SiNW-FET fabrication. a) Photomask for ohmic contacts. The chip size is 1.6×1.6 cm^2^. b) Zoom of the 2×2 mm^2^ active area with 89 nanowires. The lower doped nanowire positions are indicated in blue, the contact lines in grey. Several design of arrays (linear, rectangular, triangular) are shown. c) Optical micrograph of one triangular NW-FET array with the complementary adhesive micropattern for positioning neurons. (scale bar = 40 μm).

